# Evolutionary variations in the biofilm-associated protein BslA from the genus *Bacillus*

**DOI:** 10.1101/091884

**Authors:** Ryan J. Morris, Marieke Schor, Rachel M.C. Gillespie, Ana Sofia Ferreira, Keith M. Bromley, Lucia Baldauf, Sofia Arnaouteli, Tetyana Sukhodub, Adam Ostrowski, Laura Hobley, Chris Earl, Nicola R. Stanley-Wall, Cait E. MacPhee

## Abstract

BslA is a protein secreted by *Bacillus subtilis* which forms a hydrophobic film that coats the biofilm surface and renders it water-repellent. We have characterised three orthologues of BslA from *Bacillus amyloliquefaciens*, *Bacillus licheniformis* and *Bacillus pumilus* as well as a paralogue from *B. subtilis* called YweA. We find that the three orthologous proteins can substitute for BslA in *B. subtilis* and confer a degree of protection, whereas YweA cannot. The degree to which the proteins functionally substitute for native BslA correlates with their *in vitro* biophysical properties. Our results demonstrate the use of naturally-evolved variants to provide a framework for teasing out the molecular basis of interfacial self-assembly.

## Introduction

Many bacterial species live in multi-cellular communities known as biofilms.^1,2^ The Gram-positive bacterium *Bacillus subtilis* forms biofilms with a characteristic wrinkled morphology and a surface that is highly hydrophobic. A critical component in generating the hydrophobicity of the biofilm is the amphiphilic protein BslA.^3^ The structure of BslA contains a hydrophobic, surface-exposed ‘cap’ that sits atop a hydrophilic immunoglobulin-like domain, and is reminiscent of the fungal hydrophobins.^3–6^ We recently characterised the mechanism by which BslA remains both stable and monomeric in aqueous environments and how its adsorbs to an interface.^7^ We showed that the hydrophobic cap of BslA, in contrast to the hydrophobins, is structurally plastic; it undergoes an environmentally-responsive conformational change upon partitioning from an aqueous to a hydrophobic environment. Once at an interface, BslA self-assembles into an ordered rectangular 2D lattice that macroscopically forms an elastic protein film.^3,7,8^ Like the hydrophobins, BslA has been recognized as a ‘natural’ surface active agent that could find a range of applications in the food and personal care industries e.g. as surface modifier, coating agent, emulsifier, and/or foam stabiliser.^9–13^

Although the gross structural changes that lead to the surface-stabilising function of BslA are understood, we wished to tease out the mechanism by which BslA interacts with a hydrophilic-hydrophobic interface, i.e. what triggers the structural change between the conformational forms? Moreover, what interactions are critical for the formation of the regular 2D lattice? To address these questions, we have studied several BslA orthologues that are present in the genome of *Bacillus* species closely related to *B. subtilis*. In addition, we have investigated a paralogue of BslA called YweA. Using genetic modification and biophysical techniques, we characterize the behaviour of these variants both *in vivo* and *in vitro*. We find that key serine residues in the cap regions of the variants act as ‘switches’ that drive conformational re-arrangement at an interface. We propose a classification system for BslA and its variants based upon the behaviour of the protein films under compression. This categorization allows us to tease out the interface-protein and protein-protein interactions and suggest amino acids important for the formation of stable, highly organised 2D films.

## Results

### Protein Alignment & Choice of BslA Variants

As previously reported by Kobayashi & Iwano, a number of *Bacillus* species possess well-conserved genes that encode for homologous proteins that fall into two groups: BslA-like and YweA-like.^8^ Figure 1 shows a neighborhood joining tree of the two paralogues YweA and BslA from various *Bacillus* species. The paralogue YweA shares 67% sequence similarity with BslA and is distinguished from BslA by its lack of an N-terminal domain after the signal sequence in addition to the absence of 10 amino acids at the C-terminus that contain a conserved ‘CxC’ motif (Fig. 2A). In this work we have chosen to study the BslA orthologues produced by three species, *Bacillus amyloliquefaciens* (Ba_BslA), *Bacillus licheniformis* (Bl_BslA), and *Bacillus pumilus* (Bp_BslA), as well as the BslA paralogue YweA from *B. subtilis*. The amino acid sequences of these proteins share a high degree of similiarity to *B. subtilis* BslA (Bs_BslA), while still possessing variations that may influence their behaviour both *in vivo* and *in vitro* (Fig. 2A).

**Figure 1:**
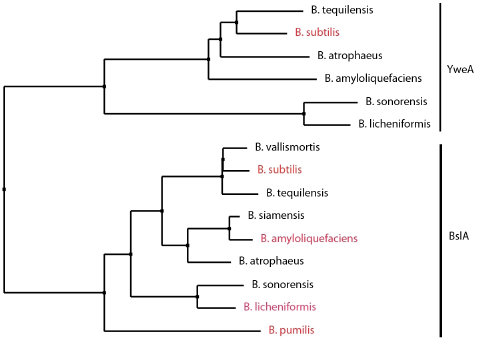
Neighbourhood joining tree of *Bacillus* species. Neighbourhood joining tree showing both YweA and BslA orthologues in other *Bacillus* species. Highlighted in red are the species that are studied in this paper.

**Figure 2:**
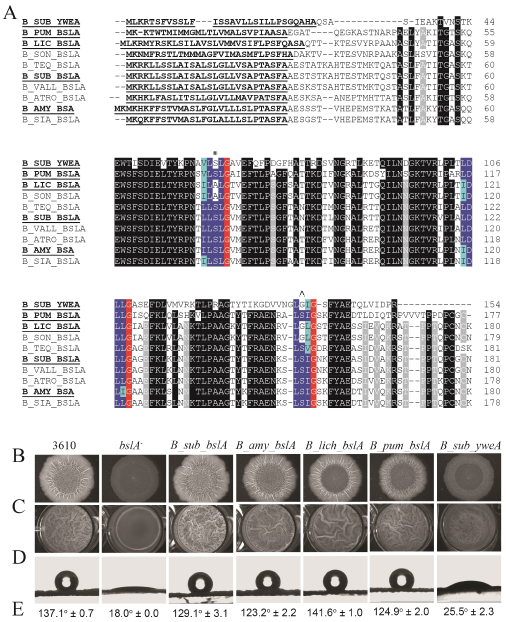
Assessing biofilm formation of heterologously expressed *bslA* variants in B. *subtilis*. (A) Amino acid alignment of BslA variants found in other *Bacillus* species. Abbreviations used are as follows: B_teq, *Bacillus tequilensis;* B_sub, *Bacillus subtilis*; B_pum, *Bacillus pumilis;* B_amy, *Bacillus amyloliquefaciens*; B_lic, *Bacillus licheniformis*; B_son, *Bacillus sonorensis*; B_teq, *Bacillus tequilensis;* B_vall, *Bacillus vallismortis;* B_atro, *Bacillus atrophaeus;* B_sia, *Bacillus siamensis*. The variants that are bolded and underlined were studied in this work. Underlined and bolded amino acids signify the signal sequence, black represents ≥90% sequence conservation, and grey 80%. Blue amino acids represent the hydrophobic cap regions, where dark blue are conserved amino acids and light blue are conservative substitutions. The red highlighted amino acids are glycines which are conserved across all species and always follow the amino acids comprising the caps. Note that YweA is differentiated from Bs_BslA and the BslA orthologues by the fact it lacks both the N-terminal region following the signalling sequence and the C-terminal domain. The * and Λ symbols indicate the cap regions containing serine residues. Biofilm phenotypes were characterized by assessing (B) complex colony morphology; (C) pellicle formation; (D,E) colony surface hydrophobicity (see Table S3 for details on strains).

### In vivo characterization of BslA variants

First, we tested whether the orthologous genes, and therefore proteins, could substitute for Bs_BslA *in vivo* in *B. subtilis*. This was achieved using heterologous expression of *bslA* variants in *B. subtilis* and biofilm formation was assessed by four criteria: complex colony morphology, pellicle formation, colony surface hydrophobicity, and sporulation ability. Wild-type *B. subtilis* strain NCIB3610 produces characteristic biofilm and pellicle morphologies with a pronounced surface hydrophobicity, reflected in the high contact angle a water droplet makes with respect to the biofilm surface (Fig. 2D, E left column). The biofilms heterologously expressing Ba_BslA showed the greatest similarity to wild-type or Bs_BslA complemented biofilms, with a characteristic wrinkled morphology and pronounced surface hydrophobicity. The *B. subtilis* biofilms substituted with Bl_BslA were also highly hydrophobic and wrinkled, however, the wrinkles in the central ‘disc’ of the colony were less pronounced and the extent of wrinkling in the pellicle was reduced. For Bp_BslA, the colony surface showed marked hydrophobicity, however colony growth appeared diminished relative to NCIB3610 and the *bslA* mutant genetically complemented with Bs_BslA. Moreover, like the *bslA* mutant genetically complemented with Bl_BslA, the wrinkles in the central ‘disc’ of the colony were less pronounced. Finally, attempts to genetically complement the *bslA* mutant with the *yweA* coding region did not reinstate either the characteristic wrinkled morphology or surface hydrophobicity of biofilms, consistent with YweA having only a minor role in the biofilm architecture (Fig. 3 and^8^).

**Figure 3:**
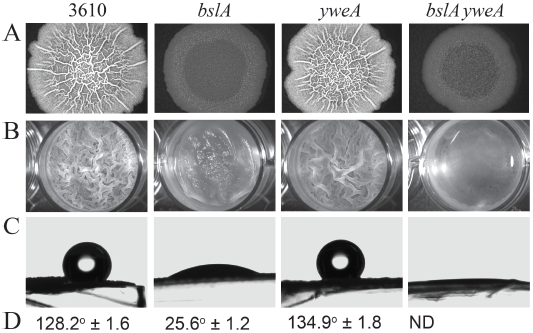
Characterization of biofilm phenotypes of *yweA* mutants. Biofilm phenotypes were characterized by assessing A, complex colony morphology; B, pellicle formation; C,D colony surface hydrophobicity for *B. subtilis* wild-type strain 3610, a *bslA* mutant, a *yweA* mutant, and a double mutant *bslA yweA* (see Table S3). Row C shows a representative sessile drop resting atop a biofilm formed from one of the strains used above. Row D are the average contact angles; ND means that a contact angle could not be determined from the images.

The ability to sporulate is a sign of a mature biofilm matrix environment.^17^ We utilized flow cytometry in conjunction with a P_*sspB-yfp*_ transcriptional reporter fusion to assess the degree of sporulation within each of the heterologously complemented biofilms. All heterologously expressed BslA orthologues were found to confer high levels of sporulation relative to a negative control, albeit not to the degree exhibited by the wild-type strain (Table 1). Taken together, these data indicate that the orthologous proteins Ba_BslA, Bl_BslA, and Bp_BslA can at least partially fulfil the role of Bs_BslA within the *B. subtilis* biofilm, largely reinstating surface hydrophobicity and gross morphological characteristics. It is notable, however, that Bl_BslA and, especially, Bp_BslA do exhibit some changes in biofilm morphology. YweA, in contrast, does not recover wild-type biofilm morphology or hydrophobicity (Figs. 2 & 3). Nonetheless, YweA does appear to contribute to the surface hydrophobicity, exacerbating the impact of the *bslA* deletion on biofilm formation (Fig. 3).

**Table 1:** Sporulation frequency assessed by flow cytometry.

| Strain | % Sporulation positive<sup>a</sup> |  |
| --- | --- | --- |
| | 0 μM IPTG | 25 μM IPTG |
| 3610 *PsspB-yfp* | 26.6 | 24.8 ± 1.63 |
| *bslA PsspB-yfp* | 5.9 ± 0.6 | 4.9 ± 0.8 |
| *bslA P<sub>IPTG</sub>-bslA<sub>B.sub</sub> PsspB-yfp* | 3.2 ± 0.2 | 14.1 ± 2.3 |
| *bslA P<sub>IPTG</sub>-bslA<sub>B.lic</sub> PsspB-yfp* | 5.6 ± 0.3 | 15.6 ± 1.2 |
| *bslA P<sub>IPTG</sub>-bslA<sub>B.amy</sub> PsspB-yfp* | 5.6 ± 0.4 | 13.0 ± 3.7 |
| *bslA P<sub>IPTG</sub>-bslA<sub>B.pum</sub> PsspB-yfp* | 5.2 ± 0.2 | 15.6 ± 1.2 |

<sup>a</sup> The percentage sporulation was calculated using fluorescence from the *PsspB-yfp* fusion as a proxy after 72 hours incubated at 30°C. NCIB3610 was used as the non-fluorescent control. For the full genotypes of the strains see Table S3. For each condition at least two independent samples were analysed. Expression of the variant gene was achieved using 25 $\mu$ M IPTG. The value presented is the mean and the error is the standard deviation.

### Biophysical assessment of protein behaviour *in vitro*

Having studied the BslA variants *in vivo*, we next characterised their behaviour in isolation using *in vitro* biophysical techniques. BslA variant proteins were expressed using standard techniques (see Experimental Procedures). The resulting mature regions of the purified proteins were first analyzed using size exclusion chromatography and SDS-PAGE (Fig. 4). As demonstrated in previous work,^3^ purified Bs_BslA is primarily composed of monomers and dimers, with a small sub-population of higher order oligomeric species (Fig. 4A, F). The orthologous BslA proteins showed similar monomer/dimer fractions as Bs-BslA, but higher order oligomers were absent. YweA, in contrast, is almost entirely monomeric (Fig. 4E).

**Figure 4:**
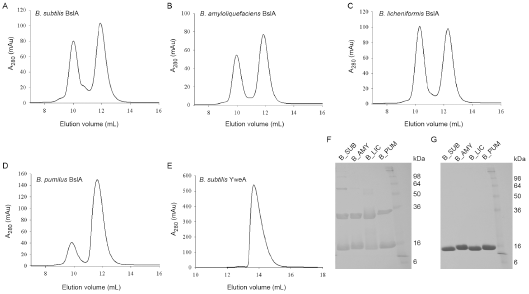
Size exclusion chromatography and SDS-PAGE analysis of Monomer-Dimer Formation. The *bslA* variant genes were cloned, overexpressed as a GST fusion and then purified. (A) Using size exclusion chromatography Bs_BslA was found to contain two primary peaks corresponding to monomer and dimer species, although there are higher-order additional peaks present. (B) Ba_BslA, (C) Bl_BslA, and (D) Bp_BslA all eluted in primarily two peaks. (E) YweA, in contrast, eluted in effectively one monomer peak. (F) SDS-PAGE analysis of recombinant protein without *β*-mercaptoethanol and (G) with *β*-mercaptoethanol.

#### Kinetics of interfacial self-assembly of BslA variants

After protein purification, we assessed the interfacial activity of the BslA variants using pendant drop tensiometry. The dynamics of interfacial protein adsorption can often be characterized by three kinetic regimes: Regime I is a ‘lag time’ where there is no apparent change in the interfacial tension, Regime II is when a sufficient proportion of protein adsorbs to the interface to produce a decrease in interfacial tension, and Regime III is reached when the interfacial tension plateaus to a roughly constant final value.^23^ Importantly, it should be stressed that once an elastic film has formed there is no longer an interfacial tension between the vapour and liquid phases and the concept of an ‘interfacial tension’ no longer applies. A good indication of when a film has formed is an increase in the error in the fit of the Young-Laplace equation to the drop shape.^24^ Bromley *et al*. showed that the only sensible data that can be extracted from interfacial tension measurements of Bs_BslA was the time at which the interfacial tension transitions between Regime I and Regime II (the ‘Regime I time’), since at any time after this transition an elastic film is present on the droplet surface.^7^ Regime I times were determined in one of two ways: (1) the transition time between regimes I and II when the fit error was still low (< 0.4 *μ*m) or (2) when the fit error increased to a threshold value (> 0.75 *μ*m). Criterion (1) or (2) was chosen by whichever occurred first. We found that the Regime I times of the orthologues Ba_BslA and Bp_BslA were within error of Bs_BslA (Fig. 5A). In contrast, the Regime I time of Bl_BslA was nearly twice as long as those of the other orthologues while the Regime I time for YweA was faster than that of Bs_BslA by ∼ 25%.

**Figure 5:**
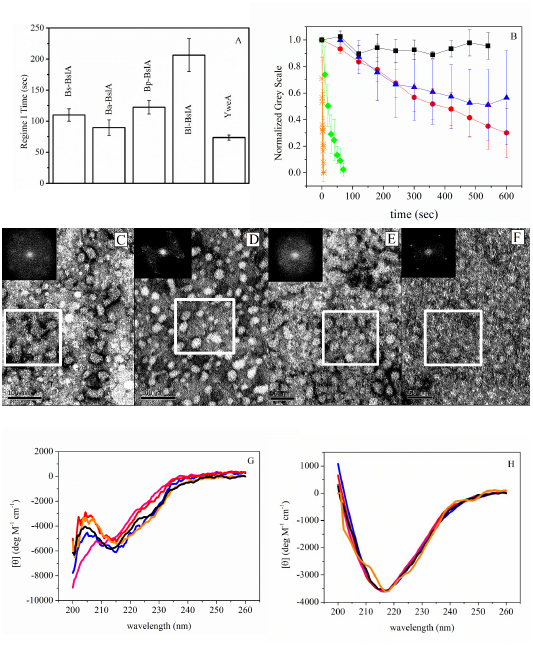
Biophysical characterization of BslA orthologues. (A) Regime I times were measured using pendant drop tensiometry. Plotted is the mean of 4 separate repeat experiments and the error bars correspond to the standard deviation. (B) Wrinkles were formed in the protein films by compressing the pendant droplet via removal of fluid. The relaxation of these wrinkles are plotted as a function of time: Bs_BslA(black squares); Ba_BslA (blue triangles), Bl_BslA (red circles); Bp_BslA (green diamonds); YweA (orange stars). Error bars represent standard deviation in relaxation times for at least N=10 separate wrinkles across the drop. (C) Inset in each image is the FFT contained within the white box. Ba_BslA visually showed ordered domains, but was not widespread.The FFT displays weak peaks indicating some ordering. (D) Bl_BslA has perceptibly larger domains of order. The FFT contains obvious peaks indicating ordering of the protein. (E) TEM of Bp_BslA films showed very weak ordering with patchy organization, which is reflected in the isotropic FFT. (F) YweA formed the ordered films that were the most similar to Bs_BslA, as can be seen from the FFT. (G) Solution state circular dichroism spectra of BslA variants and (H) circular dichroism spectra of RIMEs: Bs_BslA (black); Ba_BslA (blue); Bl_BslA (red); Bp_BslA (green); YweA (orange). Note the RIME CD spectra were normalized assuming all protein is adsorbed to emulsion droplets.

#### Relaxation dynamics of elastic films

The formation of an elastic film at a hydrophobic/hydrophilic interface is demonstrated by the observation of wrinkles following compression of the droplet (achieved by withdrawing a small volume of fluid following equilibration). One way of characterizing these films is by monitoring the formation and subsequent disappearance of these wrinkles over time, reflecting relaxation of the interfacial protein film (Fig. 5B). We have assumed that wrinkle relaxation is due to the loss of protein into the subphase, although rearrangement of the protein at the interface cannot be ruled out. For these experiments an oil/water interface was used as the small difference in density between the two phases results in rounder droplets, allowing for better imaging of the wrinkles that form. As found previously,^7^ wrinkles formed under compression in a Bs_BslA film do not relax over the timescale of the experiment. In contrast, Ba_BslA and Bl_BslA exhibit very slow relaxation, and Bp_BslA relaxes within a minute. Strikingly, relaxation of YweA is extremely rapid, occurring immediately upon compression.

#### Film organisation of BslA variants

Bs_BslA forms highly ordered 2D rectangular lattices at an air/water interface.^7^ We investigated the structure of the films formed by BslA variants using transmission electron microscopy (TEM). Protein lattices were observed for all variants, but exhibited varying degrees of order. The films formed by Ba_ BslA showed domains of ordered protein, but these were not widespread (Fig. 5C). As shown by the Fast Fourier Transform (FFT), weak peaks can be observed, indicating some level of ordering (Fig. 5C). The Bl_BslA films exhibited larger domains of ordered protein, and the FFT shows clear peaks (Fig. 5D). In contrast, Bp_BslA showed very weak ordering with only very small patches of organized structure, similar to those previously observed for the BslA L77K mutant^3^ (Fig. 5E). Thus amongst BslA orthologues, there appears to be some correlation between wrinkle relaxation and microscopic ordering: Bs_BslA films are highly ordered and wrinkles formed under compression do not relax, Ba_BslA and Bl_BslA show some degree of ordering and display slow wrinkle relaxation dynamics, whilst Bp_BslA films are not well ordered and wrinkles formed upon compression relax rapidly.

The paralogue YweA, however, does not conform to the trend observed for BslA orthologues. Indeed, YweA formed clearly defined and well ordered films similar to those formed by Bs_BslA, despite its rapid relaxation (Fig. 5B, F), calling into question the relationship between the microscopic organization of the protein within a film and film robustness.

#### CD spectroscopy of solution and interfacial states

Finally, circular dichroism (CD) spectroscopy was used to study the conformation of the BslA variants in aqueous solution and at an oil/water interface (Fig. 5G). Qualitatively, the spectra of the different BslA variants in aqueous solution are similar, with only Bp_BslA showing the loss of a feature between 210 and 218 nm (Fig. 5G). When bound to refractive indexed matched emulsions (RIMEs; Fig. 5H) all of the proteins underwent a structural transition consistent with an increase in *β*-sheet structure, reflecting rearrangement of the hydrophobic cap upon insertion into the oil phase.

### Effects of Bl_BslA cap mutations on the properties of *in vitro* protein films and *in* vivo biofilm morphology

Pendant drop tensiometry showed that it takes Bl_BslA approximately twice as long as the other orthologues to become adsorbed at an interface (Fig. 5A). This slower-than-diffusion adsorption time has previously been interpreted in relation to the energy barrier between solution and interfacial conformations of BslA,^7^ implying that there is a larger energy barrier that must be overcome for Bl_BslA adsorption. Comparison of the amino acid sequence of Bs_BslA with that of Bl_BslA reveals two amino acid differences in the hydrophobic cap region (Fig. 2A). Specifically, the first difference resides in ‘Cap 1’ (residues L76, S77, and L78 of Bs_BslA) in which serine-77 in Bs_BslA is replaced by an alanine in Bl_BslA. The second difference is in ‘Cap 3’ (residues L152, S153, and L154 of Bs_BslA) in which S153 is replaced by a glycine (G153). In contrast, these two serine resiudes are conserved in Ba_BslA and Bp_BslA, which have similar adsorption kinetics to the Bs_BslA. To investigate the role of these residues in the kinetics of adsorption and the properties of the resultant film, we produced two single amino acid mutants of Bl_BslA (A77S and G153S) and the double mutant A77S/G153S in which the cap region is identical to that of Bs_BslA. If these cap regions, and their re-orientation at an interface, determine the kinetics of adsorption, one would predict that the Regime I time of Bl_BslA(A77S/G153S) would coincide with those of Bs_BslA and the other orthologues. This is in fact what is observed as shown in Fig. 6B. In contrast, both single mutants result in Regime I times that are shorter than those of Bs_BslA and Bl_BslA(A77S/G153S) but very similar to that of YweA. Interestingly, the ‘Cap 3’ region of YweA possesses the same S → G substitution as Bl_BslA (Fig. 2A). The fact that the single mutants are more ‘YweA-like’ in terms of adsorption indicates that residues 77 and 153 help shape the energetic landscape of cap re-organization at an interface. To further explore this hypothesis, we also studied the relaxation of the films formed by these mutant proteins (Fig. 6A). Notably, wrinkles formed under compression of a Bl_BslA(A77S/G153S) droplet did not relax over the time scale of the experiment, unlike Bl_BslA which relaxes slowly, lending further support to a role for these cap residues in determining the stability and mechanical properties of the film. Consistent with this idea, the single cap mutants more closely recapitulate YweA-like behaviour, with films formed by these proteins exhibiting more rapid relaxation than both Bl_BslA and Bl_BslA(A77S/G153S) films. Indeed, films formed by Bl_BslA(A77S), the amino acid composition of which most closely resembles that of YweA, demonstrates the fastest relaxation rate of the cap mutants investigated. Intriguingly, the relaxation of wrinkles formed by this mutant appears to plateau to a non-zero value over the course of the experiment, indicating that wrinkles persist. We conclude from these results that the serine/glycine residue found in Cap 3 of BslA variants plays a prominent role in both the energetics of adsorption and the robustness of the resultant film.

**Figure 6:**
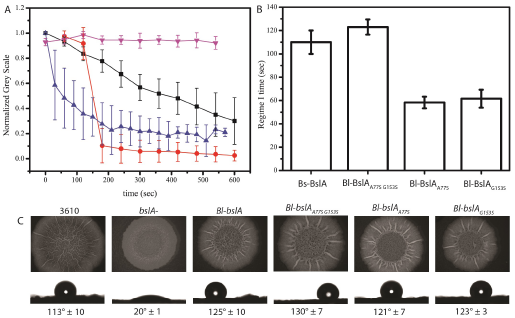
*In vitro* and *in vivo* effects of the Bl_BslA cap mutations. (A) Wrinkle relaxation measurements for Bl_BslA (black squares), Bl_BslA(A77S) (red circles), Bl_BslA(G153S) (blue triangles), Bl_BslA(A77S G153S) (upside down pink triangles). Error bars represent standard deviation in relaxation times for at least N=10 separate wrinkles across the drop. (B) Regime I times of Bs_BslA (repeated from Fig. 3 for clarity) and the Bl_BslA single and double mutants. (C) Heterologous expression of Bl_ BslA variants shows no deleterious effects on biofilm morphology or hydrophobicity relative to both wild-type Bs_BslA (as determined using the parental NCIB3610 strain (3610)) and parental Bl_BslA constructs.

Finally, we tested how the Bl_BslA cap mutations affect the structure and properties of the *B. subtilis* biofilm. When the genes encoding the Bl_BslA cap variants are heterologously expressed, the resultant biofilm morphology and hydrophobicity are indistinguishable from both wild-type Bs_BslA (as determined using the parental NCIB3610 strain) and parental Bl_BslA constructs (Fig 6C). Thus, while the *in vitro* films produced from these mutants show distinctive physical characteristics, we do not observe any differences in *in vivo* phenotype.

## Discussion

While the structural metamorphosis of the hydrophobic cap has been identified as the mechanism that allows BslA to be stable in both aqueous and hydrophobic environments, many questions remain regarding how BslA self-assembles into 2D ordered lattices and what the key amino acids are that contribute to network formation at interfaces. We have previously demonstrated that single amino acid mutations can have marked consequences for both the kinetics and energetics of adsorption along with the spatial ordering of the protein at the interface.^7,25^ Instead of using a brute-force site-directed mutagenesis approach to further illuminate the biophysical properties of BslA at an interface, we have investigated the BslA variants found in three other *Bacillus* species, along with the *B. subtilis* BslA paralogue YweA. The aim of this strategy is to lead to more efficient targeting of important amino acids for future systematic mutagenesis studies of Bs_BslA. Such knowledge could be exploited to tailor both the kinetics of interfacial adsorption as well as the mechanical properties of the protein film for the application of BslA in multiphase formulations.

We have confirmed that YweA plays no obvious role in overall biofilm formation and morphology *in vivo*, as was demonstrated in previous studies.^8^ However, the phenotype of the double deletion mutant *bslA yweA* indicates that YweA is indeed present within the biofilm and makes a small but noticeable contribution to surface hydrophobicity^8^ (Fig. 3). The major difference between YweA and the other BslA variants is the lack of both the N-and C-terminal domains. Since YweA cannot recover the characteristics of the wild-type biofilm phenotype, it is possible that the lack of one or both of these domains is functionally important for biofilm formation, in either mediating interactions between other BslA proteins, or interactions with other protein or polysaccharide components of the biofilm matrix.

Importantly, the biophysical characterization of YweA interfacial activity and self-assembly provides new insights into the relationship between the microscopic organization of the protein at the interface and resulting macroscopic properties of the surface layer. YweA interfacial films showed extremely rapid relaxation under applied compression (Fig. 5B), yet YweA films imaged by TEM showed the greatest order of all the variants (Fig. 5F). In previous studies, the cap mutant Bs_BslA(L77K) also formed films that relaxed within very short time scales. However, this variant demonstrated significant 2D disorder,^3,7^ and we suggested that the single mutation in the cap disrupted the ability of Bs_BslA(L77K) to form a space-spanning network by altering the orientation of the protein at an interface.^25^ The behaviour of YweA runs counter to this hypothesis, indicating that film order does not necessarily correlate with film robustness. It can be concluded, then, that YweA has the ability to form interfacial protein-protein lateral interactions but that these are weak, particularly compared to those formed by Bs_BslA.

As Bl_BslA differed markedly in its adsorption kinetics relative to the other BslA variants investigated here, this orthologue formed the focus of further analyses. The amino acids of Bl_BslA differ from Bs_ BslA and the other variants by two non-conservative substitutions in Cap 1 (S→A) and Cap 3 (S→G). Examination of the interfacial conformation of BslA reveals the two serines to be immediately adjacent to one another, both orientated towards the interior of protein (Fig. S1), and are likely to form a sidechain-sidechain hydrogen bond. In contrast, when the protein is in aqueous solution and the cap region is disordered, these serines are oriented outward towards the aqueous environment. The switch from an outward to an inward orientation would facilitate the conformational switch of the cap region upon contact with a hydrophobic interface, with hydrogen bonding between the residues further stabilising the interfacial conformation.

The Bl_BslA cap is more hydrophobic than Bs_BslA and the other orthologues – the polar serines are replaced by nonpolar amino acids (A77 and G153). Counter-intuitively, the higher hydrophobicity of the cap region overall does not result in faster adsorption (Fig. 5A). The resolution of this apparent paradox resides in the fact that the cap is a flexible, plastic structure. In an aqueous environment, the more hydrophobic residues of the Bl_BslA cap will most likely be oriented inwards, away from the aqueous phase, whereas in Bs_BslA the equivalent hydrophilic side chains are oriented outwards. We propose that the driving force for cap rearrangement in Bs_BslA is reorientation of these serines away from the hydrophobic interface and that this mechanism is lost in Bl_BslA, resulting in slower adsorption. Thus, we hypothesize that it is the hydrophilic residues within the cap that act as a ‘sensor’ for hydrophobic interfaces. Interaction with an apolar interface induces a reorientation of the hydrophilic amino acid(s), which in turn triggers the larger scale reorganization of the cap such that the hydrophobic residues become surface-exposed and subsequently restructure to form a three-stranded *β*-sheet.

Interestingly, the single cap mutants of Bl_BslA show shorter adsorption times, very similar to the Regime I time of YweA (Fig. 5A & Fig. 6B). YweA has a similar amino acid cap content as the Bl_ BslA(A77S) mutant, with a serine in Cap 1 and a glycine in Cap 3. For Bl_BslA(G153S), there is a hydrophobic residue in Cap 1 and a hydrophilic residue in Cap 3. We hypothesize that this combination of a hydrophilic residue in one cap strand, and a hydrophobic residue in the other, may cause a tension within the aqueous conformation of the cap as a whole or, alternatively, facilitate a rapid interconversion between states. This tension or interconversion then primes the cap to more easily undergo the conformational change when it encounters an interface, effectively lowering the energy barrier to restructuring. Moreover, when in the interfacial conformation, the remaining serine will be unfavourably partitioned into the hydrophobic protein core, but without a second serine present with which to form a hydrogen bond. This may explain why the proteins are displaced from the interface under compression.

Taking the result of this work together, we propose a categorization of BslA and its variants based upon their wrinkle relaxation behaviour:

- **Surfactant-like**: The surfactant-like proteins comprise YweA, Bp_BslA and the Bs_BslA mutants L77K and L79K.^3^ Whilst YweA differs from the other three BslA variants in its ability to form ordered lattices at an interface, wrinkles formed under compression of all four proteins relax within very short time scales. This implies that the interfacial lateral protein-protein interactions either cannot form correctly due to incorrect orientation at the interface (in the case of L77K and L79K^25^) or are weak due to substitution of critical amino acids.
- **Transiently wrinkling film formers**: The proteins that belong to this category are Ba_BslA, Bl_BslA, Bl_BslA(A77S), and Bl_BslA(G153S). In these cases, wrinkles persist for longer time scales than those formed by the surfactant-like proteins, but do eventually relax over time. Such behaviour suggests relatively weak lateral interactions between proteins at the interface, but stronger than those exhibited by the surfactant-like proteins.
- **Rigid film formers**: Bs_BslA and Bl_BslA(A77S + G153S) are members of this group. The stresses imposed via our methods are insufficient to dislodge these proteins from the interface, and moreover the entire droplet shape distorts under compression. This implies that there exist not only strong protein-interface interactions but well established protein-protein interactions within the film.

Categorizing the BslA variants in this manner allows us to identify links between the *in vivo* morphology of *Bacillus* biofilms (Figs. 3 & 4) and the interactions the proteins experience at an interface. Thus the extent to which each BslA variant rescues the wild type *B. subtilis* biofilm may be ranked and mapped to the classification system outlined above (Table 2). Bs_BslA produces rigid, elastic films *in vitro* and creates the highly wrinkled and hydrophobic biofilm phenotype associated with *B. subtilis* pellicles and colonies. Ba_BslA and Bl_BslA share similar film characteristics *in vitro* and in both cases the biofilm morphology is similar to the *B. subtilis* wild-type phenotype with some small, but discernible differences. Bp_BslA falls within the surfactant-like class and is the least effective of the orthologues at recovering the wild-type *B. subtilis* biofilm morphology. Finally, the paralogue YweA is the most surfactant-like of the proteins investigated and clearly cannot recover any features of the wild-type biofilm. Relating the biofilm morphology to the classes of film formed *in vitro* shows a clear correlation between film robustness and biofilm structure.

**Table 2:** Ranking of biofilm phenotype vs *in vitro* film classification.

| Rank | Protein | Film category |
| --- | --- | --- |
| 1 | Bs_BslA | Rigid film |
| 2 | Ba_BslA | Transient wrinkling film |
| 3 | Bl_BslA | Transient wrinkling film |
| 4 | Bp_BslA | Surfactant-like |
| 5 | YweA | Surfactant-like |

This classification provides useful insights into which residues are important for protein-protein interactions within the film. A large proportion of outward-facing, non-cap residues are conserved across all four variants. However, amino acids that are conserved between Bl_BslA and Bs_BslA but not between these two proteins and Ba_BslA and Bp_BslA are likely to affect the strength of the interaction between the proteins in the film. This suggests roles for S49, T105, Q107, N111 and E128 in the formation of the optimal lateral interactions that give rise to stable films.

Further *in vivo* investigation will be required to determine whether the biophysical differences measured as part of this study have evolved specifically to generate a fitness benefit in the distinct environmental habitat or niche occupied by each species of bacteria.

## Experimental Procedures

*Bacterial strains and growth conditions–Escherichia coli* and *Bacillus subtilis* strains used and constructed in this study are detailed in Table S1. All strains were routinely grown in Lysogeny broth (LB; 10 g NaCl, 5 g yeast extract and 10 g tryptone per litre) or on LB solidified with 1.5% (w/v) select agar (Invitrogen) at 37°C. *B. subtilis* biofilms were grown in MSgg medium (5 mM potassium phosphate and 100 mM MOPS at pH 7.0 supplemented with 2 mM MgCl_2_, 700 *μ*M CaCl_2_, 50 *μ*M MnCl_2_, 50 *μ*M FeCl3, 1 *μ*M ZnCl_2_, 2 *μ*M thiamine, 0.5% (v/v) glycerol, and 0.5% (v/v) glutamate). When required, antibiotics were used at the following concentrations: 100 *μg* ml^−1^ ampicillin, 100 *μg* ml^−1^ spectinomycin, 25 *μ*g ml^−1^ kanamycin, 1 *μ*g ml^−1^ erythromycin and 25 *μ*g ml^−1^ lincomycin. Ectopic gene expression was induced with 25 *μ*M isopropyl β-D-1 thiogalactopyranoside (IPTG). *E. coli* strain MC1061 *[F’lacIQ lacZM15 Tn10* (*tet*)] was used for the routine construction and maintenance of plasmids. *B. subtilis* 168 derivatives were generated by transformation of competent cells with plasmids using standard protocols.^14^ SPP1 phage transductions, for the introduction of DNA into *B. subtilis* strain NCIB3610, were performed as previously described.^15^

*Plasmid construction*–The primers and plasmids used in this study are presented in Tables S2 and S3 respectively. Single amino acid substitutions were generated by PCR site-directed mutagenesis using KOD Hot Start DNA Polymerase (Novagen) and the appropriate primer pairs from Table S2, with reaction conditions calculated according to the Stratagene manual for QuikChange Site-Directed Mutagenesis. The resulting mutant plasmids were digested with DpnI and transformed into competent *E. coli* MC1061 cells. All mutations were verified by DNA sequencing. For expression of BslA orthologues in *B. subtilis* under control of the IPTG-inducible Phyper-spank promoter, the *bslA* gene was amplified from genomic DNA isolated from *Bacillus licheniformis* DSM13; *Bacillus amyloliquefaciens* FZB42; and *Bacillus pumilus* SAFR-032 using the primer pairs NSW812/NSW813; NSW829/NSW830; and NSW819/NSW820 respectively. Resulting PCR products were digested with HindIII and SphI, and ligated into the *B. subtilis* shuttle vector, pDR111 (also digested HindIII/SphI), to enable subsequent gene integration into the non-essential *amyE* locus of the *B. subtilis* chromosome (final pDR111-derived plasmids are listed in Table S3). Expression of *yweA* was achieved in a similar manner. The *yweA* gene was amplified from *B. subtilis* NCIB3610 genomic DNA using primers NSW810 and NSW811, digested HindIII/SphI and ligated into HindIII/SphI-digested pDR111.

Plasmid pNW1420, for over-expression of YweA_31–155_ in *E. coli*, was generated as follows: yweA_31–155_ was amplified from *B. subtilis* NCIB3610 genomic DNA using primers NSW1853 and NSW1854 and ligated into the expression vector pET15bTEV using the NdeI and XhoI restriction sites. The resulting plasmid encodes the N-terminally His_6_-tagged fusion protein, His_6_-YweA_31–155_, which contains the recognition site of Tobacco Etch Virus (TEV) protease between His_6_ and YweA_31–155_coding regions, allowing removal of the His_6_ tag by digestion with TEV protease. Plasmids for production of BslA orthologues from *B. amyloliquefaciens*, *B. licheniformis*, and *B. pumilus* encoded truncated proteins corresponding to *B. subtilis* BslA_42–181_ (determined by sequence alignment; see Fig. 2A), on which previous analyses were based,^3,8^ and were generated as follows. For over-expression of *B. amyloliquefaciens* BslA_42–181_ (referred to herein as Ba_BslA), the appropriate DNA fragment was amplified from *B. amyloliquefaciens* FZB42 genomic DNA using primers NSW2011 and NSW2012, and ligated into the GST expression vector pGEX-6-1P via BamHI and XhoI sites. The resulting plasmid, pNW1422, encodes the N-terminally tagged fusion protein GST-Ba_BslA, from which the GST tag can be removed by TEV protease digestion. Plasmid pNW1423 for over-expression of *B. licheniformis* BslA_40–17_g (Bl_ BslA) and pNW1424 for over-expression of *B. pumilus* BslA_37–177_ (Bp_BslA), were generated in an identical manner using primer pairs NSW2013/NSW2014 and *B. licheniformis* DSM13 genomic DNA, and NSW2015/NSW2016 and *B. pumilus* SAFR-032 genomic DNA respectively. All resulting constructs were sequence-verified.

*Biofilm analysis*–Biofilm analysis was performed as previously described, with minor modifications.^15^ For complex colony formation, 10 *μ*L of the appropriate *Bacillus* culture, grown to mid-exponential phase in LB, was spotted onto MSgg medium solidified with 1.5% (w/v) agar and incubated at 30°C for 48 hours. For pellicle analysis, the LB starter culture was diluted 1:100 (v/v) in MSgg medium and incubated at 25°C for 72 hours. Both colony and pellicle biofilms were imaged using a Leica MZ16 stereoscope (Leica Microsystems).

*Colony hydrophobicity contact angle measurements*–Biofilms were grown on MSgg agar plates (as described above). A section through the mature biofilm was extracted using a scalpel and placed on a microscope slide. A 5 *μ*L droplet of sterile double-distilled water was placed on the colony using the ThetaLite Optical Tensiometer with OneAttension software. The drop was allowed to equilibrate for 5 minutes prior to imaging and contact angle measurement.

*Sporulation assays and FACS analysis*–Strains harbouring the promoter *PsspB* fused with *yfp* coding sequence were used to assess sporulation levels in mature colonies by flow cytometry.^16^ Biofilms were grown for 3 days at either 30°C or 37°C on MSgg agar without IPTG or supplemented with 25 *μ*M IPTG. After incubation, biofilms were collected, washed and fixed with 4% paraformaldehyde (PFA), as previously described.^17^ Fluorescence was analysed at the single-cell level using a BD FACSCalibur (BD Biosciences).^18^

*Protein Purification*–BslA proteins were purified as previously described,^7^ with minor changes as follows. *E. coli* BL21 (DE3) pLysS cells were transformed with the appropriate plasmids for over-expression of GST-BslA fusion proteins (described above; Table S3). For protein production, the transformed cells were grown in autoinduction medium^19^ supplemented with ampicillin (100 *μ*gmL-1) at 37°C and 200 rpm until an OD6oo of 0.9 was reached, at which point protein expression was induced by reducing the temperature to 18°C for further incubation overnight. Cells were collected by centrifugation at 4000 x *g* for 45 min and stored at −80°C until further use. For purification, bacterial pellets were thawed and resuspended in purification buffer (50 mM HEPES pH 7.5, 300 mM NaCl (for YweA), or 50 mM HEPES pH 7.5, 250 mM NaCl (for all other purifications)) supplemented with Complete EDTA-free Proteinase Inhibitors (Roche) before lysis using an Emulsiflex cell disruptor (Avestin). Unlysed cells and cell debris were removed by centrifugation at 27 000 x *g* for 20 min, and the cleared lysates incubated with Glutathione Sepharose 4B resin (GE Healthcare; for BslA) or Ni-nitrilotriacetic acid (Ni-NTA) agarose (Qiagen; for YweA) at a ratio of 750 *μ*L resin per 1 L bacterial culture with gentle rotation for 4 h at 4°C. To isolate the bound fusion proteins, the mixture of lysate plus beads was passed through a single-use 25mL gravity flow column (Bio-Rad), and the collected beads washed twice with 20 mL of the appropriate purification buffer. Untagged BslA variants were generated by resuspending the washed beads in 25 mL purification buffer supplemented with 1 mM DTT and 0.5 mg TEV protease prior to gentle rotation overnight at 4°C. To generate untagged YweA from the His6-YweA fusion protein, the overnight mixture was additionally supplemented with 250 mM imidazole. Following overnight cleavage, mixtures were again passed through gravity flow columns and the flow-through collected. To the flow-through, 750 *μ*L fresh Glutathione Sepharose plus 250 *μ*L Ni-NTA agarose (for BslA variants), or 100 *μ*L Ni-NTA agarose (for YweA), was added and the solution incubated with gentle rotation overnight at 4°C to remove the TEV protease and any unbound GST or Hisβ. The mixtures were passed through the gravity flow columns for a final time, and purified proteins collected in the flow-through. The purified proteins were then concentrated (with simultaneous exchanges into 25 mM phosphate buffer pH 7.0 where appropriate) and further purified by size exclusion chromatography (SEC) as previously described.^7^

*SDS-PAGE analysis*–SDS-PAGE analysis of BslA variants was performed using 30 *μ*g purified samples of BslA diluted 4:1 (v/v) in 4X loading buffer (6.2 g SDS, 40 mL 0.5 M Tris pH 6.8, 6.4 mL 0.1 M EDTA, 32 mL glycerol, 1 mg Bromophenol blue) either with or without β-mercaptoethanol at 4% (v/v). Samples to which β-mercaptoethanol was added were boiled at 100°C for 5 min prior to analysis, whilst samples without β-mercaptoethanol were not subjected to boiling. Proteins were run on a standard 14% polyacrylamide denaturing gel at 200V for 60 min and visualised by staining with Coomassie Blue.

*Mass spectrometry analysis*–Purified proteins were identified by excision of protein bands from SDS-PAGE gels and LC-MS-MS following tryptic digest. Protein size estimation was performed using MS-TOF and peptides identified from the MASCOT database. All procedures were performed by FingerPrints Proteomics service, School of Life Sciences, Dundee University.

*Bioinformatics*–BslA orthologues were identified using BLASTP^20,21^ using the protein sequence of BslA from *B. subtilis* as the query. BslA orthologues were distinguished from YweA based on the presence of a C-terminal region not present in YweA (,^8^ see Fig. 2A). BslA protein sequences were analysed using Jalview version 2.8.2^22^ where they were aligned using ClustalWS with the default settings. The aligned sequences were exported from Jalview into Word (Microsoft) and manually coloured for homology as indicated in the legend. A neighbourhood joining tree was generated using PAM 250 from the ClustalWS alignment in Jalview.

*Pendant drop tensiometry & and surface wrinkle relaxation quantification*–Pendant drop experiments were performed on a Krüss EasyDrop tensiometer. Protein samples were diluted in buffer and immediately placed in a syringe with a needle diameter of 1.83 mm. Images of the pendant drop are captured by a CCD camera and Krüss software fits the Young-Laplace equation to the drop shape to determine the interfacial tension. For measuring the dynamic interfacial tension, samples were prepared by diluting each protein in phosphate buffer to a concentration
 of 0.03 mg ml^−1^. Droplets were expelled in air and the interfacial tension was determined by fitting the droplet shape to the Young-Laplace equation. For the film relaxation experiments, a protein concentration of 0.2 mg ml^−1^ was used. A droplet of the protein in aqueous solution was expelled into glyceryl trioctanoate oil and allowed to equilibrate at room temperature for 30 minutes. Images were acquired at 2 fps using a digital camera. Wrinkle relaxation experiments were performed following compression of the droplet, which was achieved by retracting 5 *μ*L of the protein solution. Compression induced the formation of wrinkles in the surface layer. The wrinkles were monitored over a 10 minute period. To analyze the relaxation of the wrinkles, a line profile was drawn across the wrinkles. The line profile was plotted using the greyscale values (from 0 to 255) of each pixel along this line using the ImageJ. At least 10 wrinkles were monitored over the time course of the experiment. To plot the relaxation rate, the greyscale value of the pixels was normalized and background corrected.

*Transmission electron microscopy* (TEM)–Protein samples were deposited onto carbon-coated copper grids (Cu-grid) (TAAB Laboratories Equipment Ltd) and imaged using a Philips/FEI CM120 BioTwin transmission electron microscope. A 5 *μ*L droplet of protein (0.025 mg mL^−1^) was pipetted onto a grid and allowed to equilibrate for 5 mins before being wicked with filter paper from the side. A 5 *μ*L droplet of 2% uranyl acetate was then applied to the grid and similarly left for 5 min before being wicked from the side. The Fast Fourier Transform (FFT) of the images was performed using ImageJ software.

*Circular dichroism analysis*–Circular dichroism spectropolarimetry (CD) was performed using a Jasco J-810 spectropolarimeter. Samples were analysed at a concentration of 0.03 mgmL^1^ in a 1 cm quartz cuvette. Measurements were performed with a scan rate of 50 nm sec^−1^, a data pitch of 0.1 nm and a digital integration time of 1 sec. Twenty accumulations were measured and averaged to produce the final curve. Refractive index matched emulsions (RIMEs) were made by first preparing a 20% (v/v) decane emulsion with 0.2 mgml^−1^ of protein. The emulsion was mixed for 1 min using a rotor stator (IKA Ultra-Turrax T10) at 30,000 RPM. The emulsion was washed three times in order to remove any residual protein not adsorbed to an oil/water interface. Washes were performed by centrifuging at 1000 rpm for 20 sec, a portion of subphase was removed and replaced with buffer, then gently re-dispersed. Finally, subphase was removed and replaced with glycerol such that the final wt% of glycerol was 59% (w/v). The emulsion was then gently remixed on a rollerbank and then allowed to cream. The cream was placed in a 1 mm pathlength quartz cuvette for spectrum measurement. To prevent creaming during the experiment, the cuvette was briefly inverted between measurements to re-disperse the droplets.

## Acknowledgements

Jamie Carrington contributed a plasmid, and performed initial observations of monomeric YweA. We would like to acknowledge the Flow Cytometry and Cell Sorting Facility and the FingerPrints Proteomics Facility at the University of Dundee. This work has been supported by funding from the Engineering and Physical Sciences Research Council [EP/J007404/1] and the Biotechnology and Biological Sciences Research Council [BB/L006979/1; BB/I019464/1; BB/L006804/1].

## Author contributions

RJM designed research, performed experiments & data analysis, wrote & edited manuscript; MS performed experiments & data analysis, edited manuscript; RMCG performed experiments & data analysis, edited manuscript; KMB performed experiments & data analysis, edited manuscript; ASF provided experimental materials, performed experiments & data analysis; LB performed experiments & data analysis; SA performed experiments; TS provided experimental materials; AO performed experiments; LH performed experiments; CE performed experiments; NSW designed research, wrote & edited manuscript; CEM designed research, wrote & edited manuscript.

